# A simple, high-throughput method of protein and label removal from extracellular vesicle samples

**DOI:** 10.1101/2020.08.20.260307

**Authors:** Joshua A. Welsh, Bryce Killingsworth, Julia Kepley, Tim Traynor, Kathy McKinnon, Jason Savage, Deven Appel, Kenneth Aldape, Kevin Camphausen, Jay A. Berzofsky, Alexander R. Ivanov, Ionita H. Ghiran, Jennifer C. Jones

**Affiliations:** Translational Nanobiology Section, Laboratory of Pathology, Center for Cancer Research, National Cancer Institute, National Institutes of Health, Bethesda, MD, USA; Vaccine Branch, Center for Cancer Research, National Cancer Institute, National Institutes of Health, Bethesda, MD, USA; Laboratory of Pathology, Center for Cancer Research, National Cancer Institute, National Institutes of Health, Bethesda, MD, USA; Radiation Oncology Branch, Center for Cancer Research, National Cancer Institute, National Institutes of Health, Bethesda, MD, USA; Barnett Institute of Chemical and Biological Analysis, Department of Chemistry and Chemical Biology, Northeastern University, 360 Huntington Ave., Boston, MA, 02115, USA; Department of Medicine, Beth Israel Deaconess Medical Center, Harvard Medical School, Boston, MA, USA

## Abstract

Evidence continues to increase of the clinical utility extracellular vesicles (EVs) can provide as translational biomarkers. While a wide variety of EV isolation and purification methods have been implemented, few techniques are high-throughput and scalable for removing excess fluorescent reagents (e.g. dyes, antibodies). EVs are too small to be recovered from routine cell-processing procedures, such as filtration or centrifugation. The lack of suitable methods for removing unbound labels, especially in optical assays, is a major roadblock to accurate EV phenotyping and utilization of EV assays in a translational or clinical setting. Therefore, we developed a method for using a multi-modal resin, referred to as EV-Clean, to remove unbound labels from EV samples, and we demonstrate improvement in flow cytometric EV analysis with the use of this EV-Clean method.

## Introduction

Exosomes and ectosomes are small lipid packages released from cells, here referred to under the umbrella term of extracellular vesicles (EVs)^1, 2^. The majority of EVs have been demonstrated to be ≤100 nm in diameter, with a Power-law distribution ranging from ∼25->1000 nm^3-5^. EVs hold prospect as clinical biomarkers due to their surface and luminal cargo, hypothesized to offer a retrospective snapshot of their parent cell upon their release. Due to their small surface area, the majority of EVs typically express a very low number of copies of any one protein. Current estimates using high-sensitivity, calibrated measurements suggest the majority of EVs express ≤10 protein copies of a protein^5, 6^. This small size and limit cargo makes isolation, purification, and detection of EVs challenging.

In the most recently reported ISEV survey, which included 196 participants from 30 countries, the most reported EV isolation methods included: ultracentrifugation, density gradient, filtration, size-exclusion chromatography, precipitation, and magnetic bead capture^7, 8^. Despite, a wide variety of techniques being utilized to date for EV isolation, a gold-standard, or general consensus, is yet to emerge^9^. One of the main drawbacks of the current methodologies is their lack of high-throughput compatibility, with many techniques being labor intensive and time consuming^4, 10-12^. Isolation procedures implemented are also dependent on factors such as, the type of medium e.g. plasma, cell culture, the volume of medium e.g. µL to L, the downstream analysis technique e.g. single-particle methods or bulk methods, and the scale of isolation e.g. a couple of samples to hundreds of samples.

A wide variety of detection methods have been utilized for characterizing single EVs^5, 6, 13-18^. A common analysis technique of interest for translation studies is EV flow cytometry (EV-FC)^6, 19^. EV-FC has been utilized in a number of forms with some commercially available flow cytometers capable of detecting single-fluorescent molecules^20^. This kind of sensitivity is required to detect limited surface epitope expression due to surface area on the smallest EVs^21^. With instrumentation capable of detecting single-fluorescent molecule, it has become critical that residual or unbound fluorophore is removed from samples prior to analysis. The removal of residual or unbound fluorophore is also a highly recommended step for conventional EV-FC with lower sensitivity instrumentation as a means to increase the signal to noise ratio, and remove artefactual populations^22-24^. For this reason, the MIFlowCyt-EV reporting framework; published as a position paper to help standardize reporting of single EV flow cytometry experiments, has specific fields to demonstrate labels are not contributing or being included in EV analysis^25^. Other techniques relying on fluorescence reagents, particularly for membrane labeling, such as microscopy also require wash steps^5, 26^.

Currently, there is a gap in EV isolation and purification methods for removal of residual or unbound fluorescent labels that can be applied in a high-throughput format to small volumes, which would retain EV yield without drastically reducing sample concentration. While it is possible to titrate antibodies and fluorescent dyes to EVs for high-throughput clinical sample analysis, where samples may be in limited supply, it is neither fast, practical, or cost-effective. Here, we demonstrate EV-Clean as a simple, high-throughput method of EV purification from residual proteins and unbound fluorescent-antibodies, which can be used with µL volumes, with a limited reduction in concentration and does not fractionate EVs into several samples, that is associated with widely used size exclusion methods.

## Materials & Methods

### Blood collection & ethics

A blood samples were obtained using EDTA collection tubes. All analyses were performed in a deidentified manner, with IRB-approeved NIH intramural protocol number 02-C-0064. Plasma samples were depleted of cells and platelets by two centrifugation steps at 2500 x g at room temperature in a swing-out bucket rotor for 15 minutes with the supernatant isolated. Platelet-poor plasma samples were then stored in low-protein binding tubes (Thermo Fisher Scientific, Waltham, USA) at −80°C. Samples were thawed at 37°C for 10 minutes before being used in downstream experiments.

### Cell culture

The immature dendritic cell line DC2.4 was kindly provided by Kenneth Rock (University of Massachusetts Medical School, Boston, MA) and cultured in phenol red-free RPMI1640 medium supplemented with 10% FBS, 1% L-glutamine, 1% penicillin-streptomycin and 0.1% β-mercaptoethanol (ThermoFisher). For EV-depleted medium preparation, 20% FBS containing RPMI was ultracentrifuged for 18□hours at 100,000□*g* at 4□°C in a 45Ti fixed angle rotor using polycarbonate tubes (both from Beckman Coulter). After ultracentrifugation, the top 50□mL of medium suspension were harvested, filtered with 0.2□µm PES filter bottles and stored at 4□°C. Before using for culture, RPMI and L-glutamine, Penicillin-streptomycin and ß-mercaptoethanol were added, to achieve the concentrations before mentioned above. To produce DC2.4-derived EVs, cells were cultured for 2–3 days in EV-depleted medium and supernatants harvested before confluence was reached. Supernatants were first depleted of cells, debris and apoptotic bodies by centrifuging at 2500 g for 15 minutes twice. Supernatants were added to 100 kDa Pall Jumbosep concentrators until 5 mL of the harvested supernatant remained. 250 µL of DPBS was added to 250 µL of concentrated EVs. This 500 µL mixture was then loaded onto a qEV Original column with fractions 6-12 collected separately. Each were analysed using Nanosight and run on an SDS-PAGE gel to confirm prescience of vesicles, before fractions 8 and 9 were combined for downstream experiments.

### BSA measurements

BSA concentration were measured using a NanoDrop 2000 Spectrometer (Thermo Fisher Scientific, USA). Prior to recording concentration using the NanoDrop, the sensor was rinsed with deionized water and dried with a cotton bud before a baseline reading was taken using DPBS. 2 µL of sample was then placed on the sensor and a concentration reading was recorded three times. Recordings were exported to .xml files. Data was plotted using Prism (v8.0.1, GraphPad Software, San Diego, USA).

### SDS Page

A 10% Tris/Glycine/SDS Buffer solution was prepared with 100 mL buffer (Bio-Rad) in 900 mL tissue culture grade water. 10 µL of Bio-Rad Precision Plus ladder were added to Bio-Rad Mini-PROTEAN TGX gels (10- or 15-well). For the neat plasma samples, 15, 10, or 5 µL of plasma was added to 8.25 µL 4x Laemmli Sample Buffer (Bio-Rad, 161-0747) and 0.75 µL 55 mM 2-mercaptoethanol (Gibco, 21985). The entirety (24, 19, or 14 µL) of the sample was added to the wells of the gel. For the samples that had been previously incubated with EV-Clean, 15, 10, or 5 µL of purified sample was added to 8.25 µL 4x Laemmli Sample Buffer and 0.75 µL 55 mM 2-mercaptoethanol. The entirety (24, 19, or 14 µL) of the mixture was added to the wells of the gel. Antibody removal was tested by suspending 0.5 µg of IgG-PE-CD147 (BioLegend, Cat. 306212) and IgG-APC-CD147 (BioLegend, Cat. 306214) in a final volume of 50 µL adding to 100 µL of EV-Clean for 30 minutes. Post-incubation 20 µL of supernatant was added to 8.25 µL 4x Laemmli Sample Buffer (Bio-Rad, 161-0747), 0.75 µL 55 mM 2-mercaptoethanol (Gibco, 21985), and 20 µL of SDS buffer. The final volume of 50 µL was added to each well of a 10 well Bio-Rad Mini-PROTEAN TGX gel.

With a Mini-PROTEAN Tetra System connected to a Bio-Rad Power Pac 1000, SDS-PAGE was run at a constant voltage of 200 V until the bands ran off the bottom of the gels. The stain-free gel was activated and imaged under the Bio-Rad ChemiDoc Touch Imaging System. Gels were analysed using Image Lab software (v6.0.1, Bio-Rad, Hercules, USA)

### Nanoparticle Tracking Analysis (NTA)

Particle concentration and diameter distribution were characterized by NTA with a NanoSight LM10 instrument (Malvern, UK), equipped with a 405□nm LM12 module and EMCCD camera (DL-658-OEM-630, Andor). Video acquisition was performed with NTA software v3.2, using a camera level of 14. Three 30□second videos were captured per sample. Post-acquisition video analysis used the following settings: minimum track length = 5, detection threshold = 4, automatic blur size = 2-pass, maximum jump size = 12.0. Exported datasets were compiled and plotted using scripts written in MATLAB v9.3.0 (The MathWorks Inc., USA). Samples were diluted to have a concentration in the region of 1×10^8^ to 1×10^9^ particles mL^-1^.

### BSA removal using EV-Clean

Samples containing approximately 400 µg of BSA diluted in 75 µL DPBS were aliquoted into PCR tubes containing 100 µL of DPBS-washed CaptoCore -700 or –400 (GE Biosciences) Samples were incubated at 4 °C for 30 minutes before the top 75 µL of supernatant was then removed and added to another 100 µL of DPBS-washed EV-Clean, mixed, and incubated for a further 30 minutes at 4 °C.

### Plasma protein removal using EV-Clean

15, 10, and 5 µL of platelet-poor plasma was aliquoted into PCR tubes containing 100 µL of DPBS-washed EV-Clean. Samples were incubated at 4 °C for 30 minutes before the top 15, 10, or 5 µL of supernatant was then removed and added to another 100 µL of DPBS-washed EV-Clean, mixed, and incubated for a further 30 minutes at 4 °C.

### EV CFSE-labeling

CFSE-labelling of DC2.4 EVs was carried out as described previously^27^. Briefly, 15 µL of 1×10^9^ DC2.4 EVs, pooled from qEV original columns fraction 8 and 9, were added to 15 µL of 40 µM of CFDA-SE (Thermo Fisher Scientific). This was protected from light and incubated at 37 °C for 2 hours. 70 µL of DPBS was added to the stained sample. This was repeated for each sample. Excess dye removal using size-exclusion chromatography used NAP-5 columns, loading 100 µL of CFSE-stained EVs and collecting fractions 3 and 4 for purified EVs, having a final volume of 500 µL. For dye removal using EV-Clean, 100 µL of CFSE-stained EVs were added to 100 µL of DPBS-washed EV-Clean in a PCR tube, mixed, and incubated at 4 °C for 30 minutes. The top 100 µL of supernatant was then removed and added to another 100 µL of DPBS-washed EV-Clean, mixed, and incubated for a further 30 minutes at 4 °C. Post-dye removal all samples were transferred to 1.5 mL low protein binding tubes (Thermo Fisher Scientific) with 1×10^9^ 200 nm Red FluoSpheres (Thermo Fisher Scientific) added, before being diluted to a final volume of 1 mL for analysis by nanoFACS.

### Flow Cytometry of EVs

Flow cytometric analysis of CFSE EVs was carried out using previously published NanoFACS methodology^27^. Briefly, an Astrios EQ jet-in-air system (Beckman Coulter), configured with 5 lasers (355, 405, 488, 561 and 640□nm wavelength), where SSC can be detected and used as a trigger at laser wavelength with the exception of the 355 nm laser. EV analyses were carried out using a 561-SSC trigger with the 561-SSC voltage and threshold settings adjusted to allow ∼10,000 events of background reference noise per second. Samples were loaded and run for 5□minutes until the event rate was stable, and then recorded for 30 seconds. All samples were run at a 0.2 psi differential pressure, monitoring stability closely. Data was acquired using Summit v6 (Beckman Coulter) and analyzed with FlowJo v10.1r5 (TreeStar, USA). CFSE fluorescence data was calibrated using FITC MESF calibration beads using FCM_PASS_ software (v.3.03)^28, 29^. Full calibration details can be found in the MIFlowCyt-EV report, **Supplementary Information 1**^25^. Flow cytometric analysis of EV recovery was carried out using a Cytek Aurora (Cytek Biosciences), configured with 4 lasers (405, 488, 561, 640 nm) with a custom modified 405 nm detector. Diameter was calculated for EVs using FCM_PASS_ software (v3.03)^28^. Light scatter parameters were calibrated into full calibration details can be found in the MIFlowCyt-EV report, **Supplementary Information 2**^25^.

## Results

Optimal incubation times for protein removal were tested with bovine serum albumin (BSA). These showed that 30 minutes and 1-hour incubation had minor differences, **Figure 1**. After a single incubation with EV-Clean the BSA content dropped from ∼400 µg to ∼70 and ∼60 µg, respectively, **Figure 1**. A 4-hour single incubation showed a decrease from ∼400 µg to no detectable BSA. To determine whether a second, sequential incubation could speed up this process, the use of an additional 30-minute incubation was tested after each of the first incubations. After the second incubation no detectable BSA was observed after any of the preceding incubation times. We therefore conclude that two 30-minute incubation are sufficient as an incubation period to work from.

**Figure 1.**
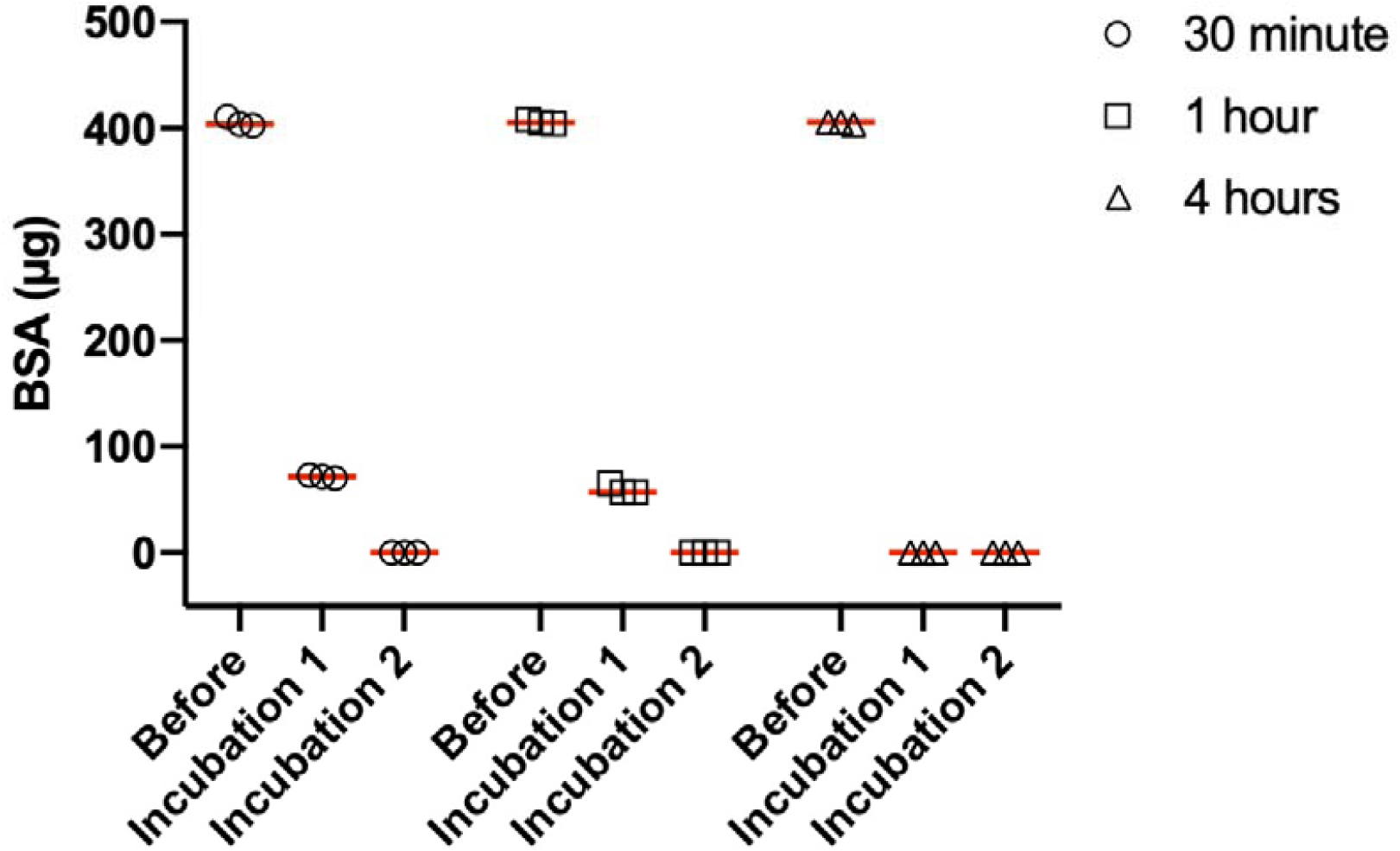
BSA removal using EV-Clean. The removal of 75 µL purified bovine serum album (BSA) using an initial incubation with 100 µL EV-Clean for either 30 minutes, 1 hour, or 4 hours followed by a 30 minute incubation with EV-clean.

The ability of EV-Clean to remove the large amount of soluble protein from a heterogeneous solution was tested using plasma samples with the effects of purification observed by SDS-PAGE, **Figure 2**. 15 µL of plasma after a single 30-minute incubation with EV-Clean shows a significantly reduced signal when compared to neat plasma. This reduction in protein content shows no observable bias in protein size with all observable protein ≦300 kDa showing depletion. A second incubation again further depletes all observable protein with only faint bands visible at ∼13, 50, 65, 185, 311 kDa. This was repeated for 10 µL and 5 µL of plasma. With 5 µL of plasma bands are only faint bands were only visible at ∼50 and 311 kDa indicating the majority of small proteins can be depleted from plasma with 100 µL of EV-Clean and two 30-minute incubations.

**Figure 2.**
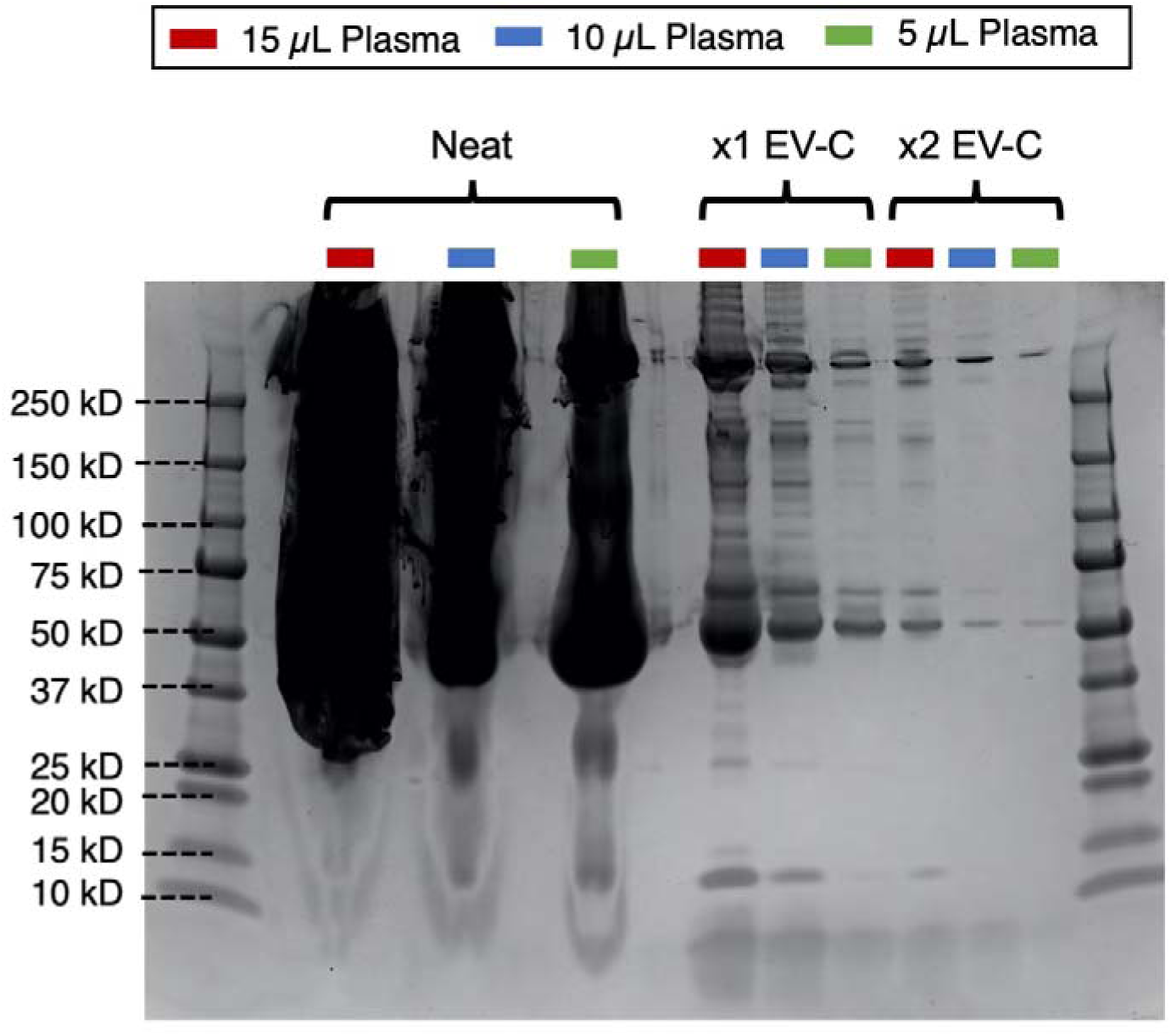
Protein removal from plasma. The ability of EV-Clean to remove protein from plasma was tested using 15 µL (red), 10 µL (blue), and 5 µL (green) of neat platelet-depleted plasma with EV-Clean for one incubation and two incubations for 30 minutes.

EV-Clean was next tested for its ability to remove fluorescently-conjugated IgG antibodies. R-phycoerythrin (R-PE) and allophycocyanin (APC) were chosen due to their large size, 250 and 105 kDa, respectively. Both R-PE-IgG1 and APC-IgG1 were significantly depleted when incubated with EV-Clean, **Figure 3A**. Along with antibodies, molecular labels are used for staining EVs. The ability of EV-Clean to remove excess CFSE from stained EV samples compared to a previously published method using size exclusion chromatography (SEC) using nanoFACS. The instrument reference noise, measured with PBS and unstained EVs, had a median brightness of 48-49 fluorescein (FITC) molecules of equivalent soluble fluorophore (MESF) units. EVs without the removal of excess CFSE label resulted in the cytometer noise being raised to a median brightness of 437 FITC MESF units, **Figure 3B**. By removing the excess CFSE, the noise level remained low with the SEC and EV-Clean purification methods having a median brightness of 58 and 54 FITC MESF units, respectively, **Figure 3B**.

**Figure 3.**
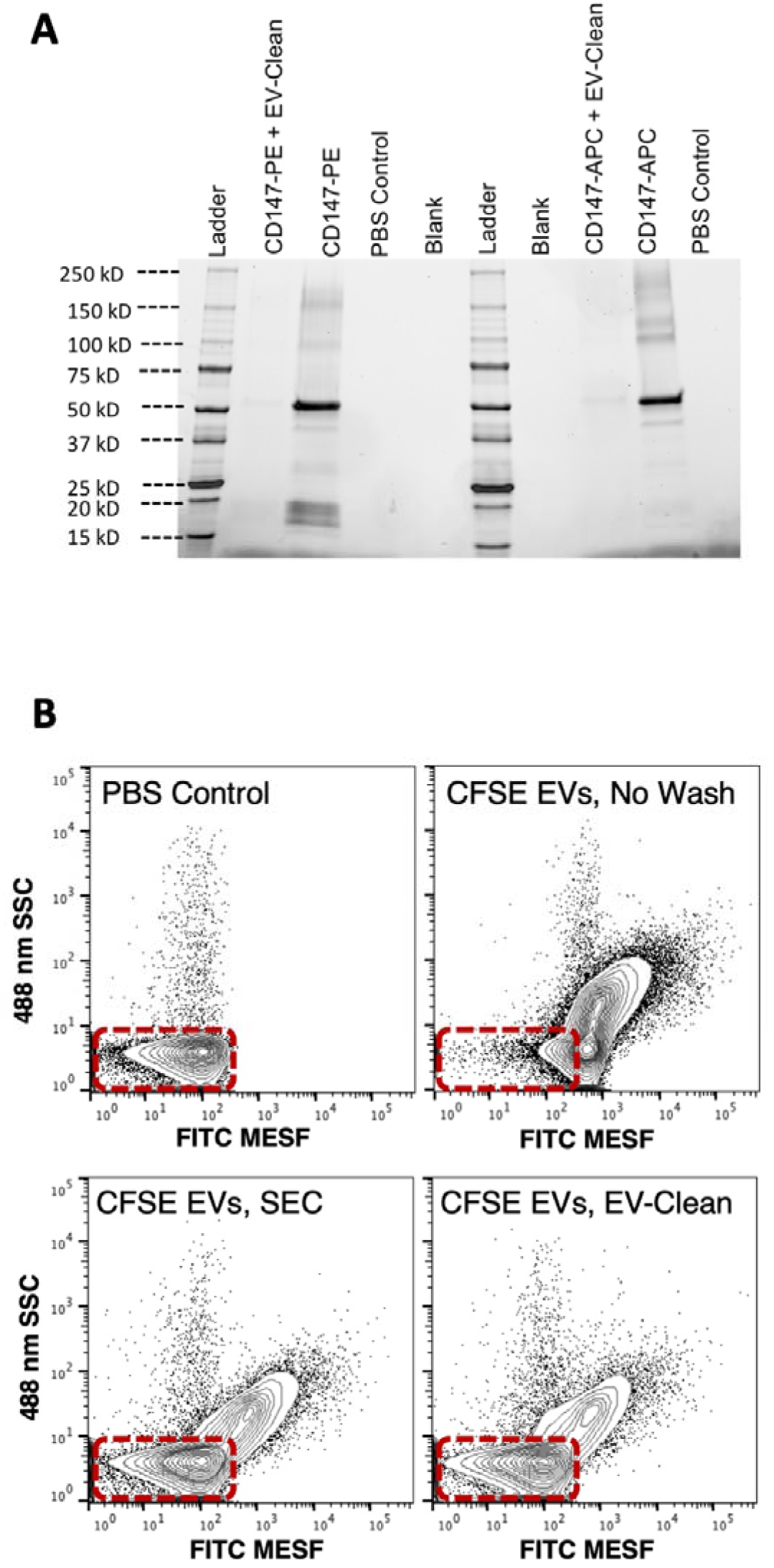
Fluorescent label removal. The ability of EV-Clean to remove fluorescent labels was tested using 1 µg of IgG antibodies conjugated to large fluorophores; phycoerythrin (PE) and allophycocyanin (APC). A comparison of EV detectability from background signal was investigated. DC2.4 EVs were detected using flow cytometry with samples stained with 20 µM of CFDA-SE with no purification (C-top right), stained and the purified used size exclusion chromatography (C-bottom left), and stained and purified using EV-Clean (C-bottom right). A buffer only control to represent the true background noise of the instrument is also shown (C-top left)

Finally, the affect of EV-Clean on the detectable EV concentration after removing unbound label was evaluated, **Figure 4**. Recovery was assessed by gating single EVs between 115 to 200 nm, **Figure 4A**. After one incubation of EV-Clean the detectable recovery was 75%, with a subsequent reduction after a second incubation to 51%. In summary, the use of EV-Clean demonstrates a 75% EV recovery after each incubalion and >95% label removal, without dilution, for each incubation.

**Figure 4.**
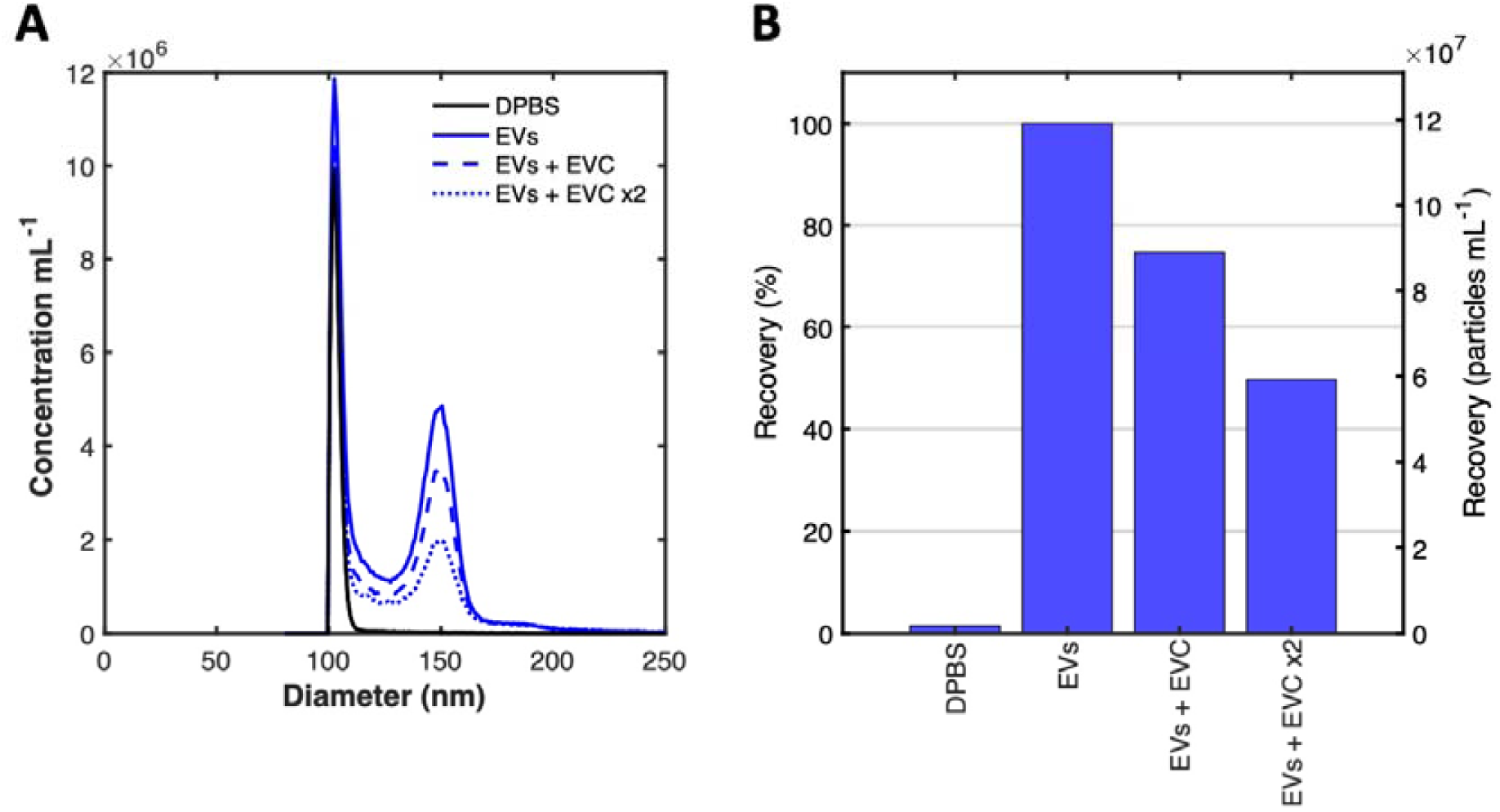
Affect of EV-clean on EV recovery. The influence of different permutation of EV-clean on detectable EV recovery was assessed using flow cytometry (A). The percentange recovery of EVs gated from 115-200 nm using EV-Clean and incubated once or twice for 30 minutes was assessed (B).

## Discussion

We have demonstrated that EV-Clean is able to significantly deplete soluble proteins from heterogeneous samples such as plasma in a form-factor that is compatible with 96-well plates and robotics. This methodology requires up to just two 30-minute incubations to achieve significant protein-depletion. Furthermore, we have demonstrated that the detectable EV concentration using flow cytometry is ∼75% using EV-clean, making it a useful tool for EV-isolation from heterogenous samples containing many soluble proteins. Due EV-clean protein removal being a multi-mode process of interculation and affinity capture, it is possible that EV recovery is higher in samples containing more proteins than the size-exclusion purified samples that were tested. We have also demonstrated the use of EV-Clean for EV-labelling is beneficial for high-sensitivity analysis techniques, such as flow cytometry, where the removal of excess label can reduce sensitivity and several samples need to be prepared simultaneously. While this proposed method offers potential for relatively small volume, high-throughput purification applications, it may be less suitable for large volume applications such as tissue culture supernatants.

Future development of EV-Clean as a reliable purification method for heterogenous samples, such as plasma, will require additions to also deplete large and abundant particles, such as lipoproteins, that could confound EV data after soluble-protein removal. Consistent packing and receptacle formats are also required to ensure consistent results, as is the case for size-exclusion chromatography methods.

## Acknowledgements

Research for this manuscript was supported by NIH U01 HL126497, NIH R01 CA218500, NIH UG3 TR002881, NIH ZIA BC011502, NIH R01 GM120272, NIH R35 GM136421, and NIH ZIA BC011503.

## References

1. Thery, C.; Witwer, K. W.; Aikawa, E.; Alcaraz, M. J.; Anderson, J. D.; Andriantsitohaina, R.; Antoniou, A.; Arab, T.; Archer, F.; Atkin-Smith, G. K.; Ayre, D. C.; Bach, J. M.; Bachurski, D.; Baharvand, H.; Balaj, L.; Baldacchino, S.; Bauer, N. N.; Baxter, A. A.; Bebawy, M.; Beckham, C.; Bedina Zavec, A.; Benmoussa, A.; Berardi, A. C.; Bergese, P.; Bielska, E.; Blenkiron, C.; Bobis-Wozowicz, S.; Boilard, E.; Boireau, W.; Bongiovanni, A.; Borras, F. E.; Bosch, S.; Boulanger, C. M.; Breakefield, X.; Breglio, A. M.; Brennan, M. A.; Brigstock, D. R.; Brisson, A.; Broekman, M. L.; Bromberg, J. F.; Bryl-Gorecka, P.; Buch, S.; Buck, A. H.; Burger, D.; Busatto, S.; Buschmann, D.; Bussolati, B.; Buzas, E. I.; Byrd, J. B.; Camussi, G.; Carter, D. R.; Caruso, S.; Chamley, L. W.; Chang, Y. T.; Chen, C.; Chen, S.; Cheng, L.; Chin, A. R.; Clayton, A.; Clerici, S. P.; Cocks, A.; Cocucci, E.; Coffey, R. J.; Cordeiro-da-Silva, A.; Couch, Y.; Coumans, F. A.; Coyle, B.; Crescitelli, R.; Criado, M. F.; D’Souza-Schorey, C.; Das, S.; Datta Chaudhuri, A.; de Candia, P.; De Santana, E. F.; De Wever, O.; Del Portillo, H. A.; Demaret, T.; Deville, S.; Devitt, A.; Dhondt, B.; Di Vizio, D.; Dieterich, L. C.; Dolo, V.; Dominguez Rubio, A. P.; Dominici, M.; Dourado, M. R.; Driedonks, T. A.; Duarte, F. V.; Duncan, H. M.; Eichenberger, R. M.; Ekstrom, K.; El Andaloussi, S.; Elie-Caille, C.; Erdbrugger, U.; Falcon-Perez, J. M.; Fatima, F.; Fish, J. E.; Flores-Bellver, M.; Forsonits, A.; Frelet-Barrand, A.; Fricke, F.; Fuhrmann, G.; Gabrielsson, S.; Gamez-Valero, A.; Gardiner, C.; Gartner, K.; Gaudin, R.; Gho, Y. S.; Giebel, B.; Gilbert, C.; Gimona, M.; Giusti, I.; Goberdhan, D. C.; Gorgens, A.; Gorski, S. M.; Greening, D. W.; Gross, J. C.; Gualerzi, A.; Gupta, G. N.; Gustafson, D.; Handberg, A.; Haraszti, R. A.; Harrison, P.; Hegyesi, H.; Hendrix, A.; Hill, A. F.; Hochberg, F. H.; Hoffmann, K. F.; Holder, B.; Holthofer, H.; Hosseinkhani, B.; Hu, G.; Huang, Y.; Huber, V.; Hunt, S.; Ibrahim, A. G.; Ikezu, T.; Inal, J. M.; Isin, M.; Ivanova, A.; Jackson, H. K.; Jacobsen, S.; Jay, S. M.; Jayachandran, M.; Jenster, G.; Jiang, L.; Johnson, S. M.; Jones, J. C.; Jong, A.; Jovanovic-Talisman, T.; Jung, S.; Kalluri, R.; Kano, S. I.; Kaur, S.; Kawamura, Y.; Keller, E. T.; Khamari, D.; Khomyakova, E.; Khvorova, A.; Kierulf, P.; Kim, K. P.; Kislinger, T.; Klingeborn, M.; Klinke, D. J., 2nd; Kornek, M.; Kosanovic, M. M.; Kovacs, A. F.; Kramer-Albers, E. M.; Krasemann, S.; Krause, M.; Kurochkin, I. V.; Kusuma, G. D.; Kuypers, S.; Laitinen, S.; Langevin, S. M.; Languino, L. R.; Lannigan, J.; Lasser, C.; Laurent, L. C.; Lavieu, G.; Lazaro-Ibanez, E.; Le Lay, S.; Lee, M. S.; Lee, Y. X. F.; Lemos, D. S.; Lenassi, M.; Leszczynska, A.; Li, I. T.; Liao, K.; Libregts, S. F.; Ligeti, E.; Lim, R.; Lim, S. K.; Line, A.; Linnemannstons, K.; Llorente, A.; Lombard, C. A.; Lorenowicz, M. J.; Lorincz, A. M.; Lotvall, J.; Lovett, J.; Lowry, M. C.; Loyer, X.; Lu, Q.; Lukomska, B.; Lunavat, T. R.; Maas, S. L.; Malhi, H.; Marcilla, A.; Mariani, J.; Mariscal, J.; Martens-Uzunova, E. S.; Martin-Jaular, L.; Martinez, M. C.; Martins, V. R.; Mathieu, M.; Mathivanan, S.; Maugeri, M.; McGinnis, L. K.; McVey, M. J.; Meckes, D. G., Jr.; Meehan, K. L.; Mertens, I.; Minciacchi, V. R.; Moller, A.; Moller Jorgensen, M.; Morales-Kastresana, A.; Morhayim, J.; Mullier, F.; Muraca, M.; Musante, L.; Mussack, V.; Muth, D. C.; Myburgh, K. H.; Najrana, T.; Nawaz, M.; Nazarenko, I.; Nejsum, P.; Neri, C.; Neri, T.; Nieuwland, R.; Nimrichter, L.; Nolan, J. P.; Nolte-’t Hoen, E. N.; Noren Hooten, N.; O’Driscoll, L.; O’Grady, T.; O’Loghlen, A.; Ochiya, T.; Olivier, M.; Ortiz, A.; Ortiz, L. A.; Osteikoetxea, X.; Ostergaard, O.; Ostrowski, M.; Park, J.; Pegtel, D. M.; Peinado, H.; Perut, F.; Pfaffl, M. W.; Phinney, D. G.; Pieters, B. C.; Pink, R. C.; Pisetsky, D. S.; Pogge von Strandmann, E.; Polakovicova, I.; Poon, I. K.; Powell, B. H.; Prada, I.; Pulliam, L.; Quesenberry, P.; Radeghieri, A.; Raffai, R. L.; Raimondo, S.; Rak, J.; Ramirez, M. I.; Raposo, G.; Rayyan, M. S.; Regev-Rudzki, N.; Ricklefs, F. L.; Robbins, P. D.; Roberts, D. D.; Rodrigues, S. C.; Rohde, E.; Rome, S.; Rouschop, K. M.; Rughetti, A.; Russell, A. E.; Saa, P.; Sahoo, S.; Salas-Huenuleo, E.; Sanchez, C.; Saugstad, J. A.; Saul, M. J.; Schiffelers, R. M.; Schneider, R.; Schoyen, T. H.; Scott, A.; Shahaj, E.; Sharma, S.; Shatnyeva, O.; Shekari, F.; Shelke, G. V.; Shetty, A. K.; Shiba, K.; Siljander, P. R.; Silva, A. M.; Skowronek, A.; Snyder, O. L., 2nd; Soares, R. P.; Sodar, B. W.; Soekmadji, C.; Sotillo, J.; Stahl, P. D.; Stoorvogel, W.; Stott, S. L.; Strasser, E. F.; Swift, S.; Tahara, H.; Tewari, M.; Timms, K.; Tiwari, S.; Tixeira, R.; Tkach, M.; Toh, W. S.; Tomasini, R.; Torrecilhas, A. C.; Tosar, J. P.; Toxavidis, V.; Urbanelli, L.; Vader, P.; van Balkom, B. W.; van der Grein, S. G.; Van Deun, J.; van Herwijnen, M. J.; Van Keuren-Jensen, K.; van Niel, G.; van Royen, M. E.; van Wijnen, A. J.; Vasconcelos, M. H.; Vechetti, I. J., Jr.; Veit, T. D.; Vella, L. J.; Velot, E.; Verweij, F. J.; Vestad, B.; Vinas, J. L.; Visnovitz, T.; Vukman, K. V.; Wahlgren, J.; Watson, D. C.; Wauben, M. H.; Weaver, A.; Webber, J. P.; Weber, V.; Wehman, A. M.; Weiss, D. J.; Welsh, J. A.; Wendt, S.; Wheelock, A. M.; Wiener, Z.; Witte, L.; Wolfram, J.; Xagorari, A.; Xander, P.; Xu, J.; Yan, X.; Yanez-Mo, M.; Yin, H.; Yuana, Y.; Zappulli, V.; Zarubova, J.; Zekas, V.; Zhang, J. Y.; Zhao, Z.; Zheng, L.; Zheutlin, A. R.; Zickler, A. M.; Zimmermann, P.; Zivkovic, A. M.; Zocco, D.; Zuba-Surma, E. K., Minimal information for studies of extracellular vesicles 2018 (MISEV2018): a position statement of the International Society for Extracellular Vesicles and update of the MISEV2014 guidelines. J Extracell Vesicles 2018, 7 (1), 1535750.

2. Lotvall, J.; Hill, A. F.; Hochberg, F.; Buzas, E. I.; Di Vizio, D.; Gardiner, C.; Gho, Y. S.; Kurochkin, I. V.; Mathivanan, S.; Quesenberry, P.; Sahoo, S.; Tahara, H.; Wauben, M. H.; Witwer, K. W.; Thery, C., Minimal experimental requirements for definition of extracellular vesicles and their functions: a position statement from the International Society for Extracellular Vesicles. J Extracell Vesicles 2014, 3, 26913.

3. van der Pol, E.; Coumans, F. A.; Grootemaat, A. E.; Gardiner, C.; Sargent, I. L.; Harrison, P.; Sturk, A.; van Leeuwen, T. G.; Nieuwland, R., Particle size distribution of exosomes and microvesicles determined by transmission electron microscopy, flow cytometry, nanoparticle tracking analysis, and resistive pulse sensing. J Thromb Haemost 2014, 12 (7), 1182–92.

4. Tian, Y.; Gong, M.; Hu, Y.; Liu, H.; Zhang, W.; Zhang, M.; Hu, X.; Aubert, D.; Zhu, S.; Wu, L.; Yan, X., Quality and efficiency assessment of six extracellular vesicle isolation methods by nano-flow cytometry. J Extracell Vesicles 2020, 9 (1), 1697028.

5. Lennon, K. M.; Wakefield, D. L.; Maddox, A. L.; Brehove, M. S.; Willner, A. N.; Garcia-Mansfield, K.; Meechoovet, B.; Reiman, R.; Hutchins, E.; Miller, M. M.; Goel, A.; Pirrotte, P.; Van Keuren-Jensen, K.; Jovanovic-Talisman, T., Single molecule characterization of individual extracellular vesicles from pancreatic cancer. J Extracell Vesicles 2019, 8 (1), 1685634.

6. Tian, Y.; Ma, L.; Gong, M.; Su, G.; Zhu, S.; Zhang, W.; Wang, S.; Li, Z.; Chen, C.; Li, L.; Wu, L.; Yan, X., Protein Profiling and Sizing of Extracellular Vesicles from Colorectal Cancer Patients via Flow Cytometry. Acs Nano 2018, 12 (1), 671–680.

7. Gardiner, C.; Di Vizio, D.; Sahoo, S.; Thery, C.; Witwer, K. W.; Wauben, M.; Hill, A. F., Techniques used for the isolation and characterization of extracellular vesicles: results of a worldwide survey. Journal of Extracellular Vesicles 2016, 5.

8. Consortium, E.-T.; Van Deun, J.; Mestdagh, P.; Agostinis, P.; Akay, O.; Anand, S.; Anckaert, J.; Martinez, Z. A.; Baetens, T.; Beghein, E.; Bertier, L.; Berx, G.; Boere, J.; Boukouris, S.; Bremer, M.; Buschmann, D.; Byrd, J. B.; Casert, C.; Cheng, L.; Cmoch, A.; Daveloose, D.; De Smedt, E.; Demirsoy, S.; Depoorter, V.; Dhondt, B.; Driedonks, T. A.; Dudek, A.; Elsharawy, A.; Floris, I.; Foers, A. D.; Gartner, K.; Garg, A. D.; Geeurickx, E.; Gettemans, J.; Ghazavi, F.; Giebel, B.; Kormelink, T. G.; Hancock, G.; Helsmoortel, H.; Hill, A. F.; Hyenne, V.; Kalra, H.; Kim, D.; Kowal, J.; Kraemer, S.; Leidinger, P.; Leonelli, C.; Liang, Y.; Lippens, L.; Liu, S.; Lo Cicero, A.; Martin, S.; Mathivanan, S.; Mathiyalagan, P.; Matusek, T.; Milani, G.; Monguio-Tortajada, M.; Mus, L. M.; Muth, D. C.; Nemeth, A.; Nolte-’t Hoen, E. N.; O’Driscoll, L.; Palmulli, R.; Pfaffl, M. W.; Primdal-Bengtson, B.; Romano, E.; Rousseau, Q.; Sahoo, S.; Sampaio, N.; Samuel, M.; Scicluna, B.; Soen, B.; Steels, A.; Swinnen, J. V.; Takatalo, M.; Thaminy, S.; Thery, C.; Tulkens, J.; Van Audenhove, I.; van der Grein, S.; Van Goethem, A.; van Herwijnen, M. J.; Van Niel, G.; Van Roy, N.; Van Vliet, A. R.; Vandamme, N.; Vanhauwaert, S.; Vergauwen, G.; Verweij, F.; Wallaert, A.; Wauben, M.; Witwer, K. W.; Zonneveld, M. I.; De Wever, O.; Vandesompele, J.; Hendrix, A., EV-TRACK: transparent reporting and centralizing knowledge in extracellular vesicle research. Nat Methods 2017, 14 (3), 228–232.

9. Witwer, K. W.; Buzas, E. I.; Bemis, L. T.; Bora, A.; Lasser, C.; Lotvall, J.; Nolte-’t Hoen, E. N.; Piper, M. G.; Sivaraman, S.; Skog, J.; Thery, C.; Wauben, M. H.; Hochberg, F., Standardization of sample collection, isolation and analysis methods in extracellular vesicle research. J Extracell Vesicles 2013, 2.

10. Jeurissen, S.; Vergauwen, G.; Van Deun, J.; Lapeire, L.; Depoorter, V.; Miinalainen, I.; Sormunen, R.; Van den Broecke, R.; Braems, G.; Cocquyt, V.; Denys, H.; Hendrix, A., The isolation of morphologically intact and biologically active extracellular vesicles from the secretome of cancer-associated adipose tissue. Cell Adh Migr 2017, 11 (2), 196–204.

11. Takov, K.; Yellon, D. M.; Davidson, S. M., Comparison of small extracellular vesicles isolated from plasma by ultracentrifugation or size-exclusion chromatography: yield, purity and functional potential. J Extracell Vesicles 2019, 8 (1), 1560809.

12. Witwer, K. W.; Buzas, E. I.; Bemis, L. T.; Bora, A.; Lasser, C.; Lotvall, J.; Nolte-’t Hoen, E. N.; Piper, M. G.; Sivaraman, S.; Skog, J.; Thery, C.; Wauben, M. H.; Hochberg, F., Standardization of sample collection, isolation and analysis methods in extracellular vesicle research. J Extracell Vesicles 2013, 2 (1), 20360.

13. Issadore, D.; Min, C.; Liong, M.; Chung, J.; Weissleder, R.; Lee, H., Miniature magnetic resonance system for point-of-care diagnostics. Lab on a chip 2011, 11 (13), 2282–7.

14. Shao, H.; Chung, J.; Balaj, L.; Charest, A.; Bigner, D. D.; Carter, B. S.; Hochberg, F. H.; Breakefield, X. O.; Weissleder, R.; Lee, H., Protein typing of circulating microvesicles allows real-time monitoring of glioblastoma therapy. Nat Med 2012, 18 (12), 1835–40.

15. Shao, H.; Yoon, T. J.; Liong, M.; Weissleder, R.; Lee, H., Magnetic nanoparticles for biomedical NMR-based diagnostics. Beilstein J Nanotech 2010, 1, 142–54.

16. van der Pol, E.; Coumans, F.; Varga, Z.; Krumrey, M.; Nieuwland, R., Innovation in detection of microparticles and exosomes. J Thromb Haemost 2013, 11 Suppl 1, 36–45.

17. van der Pol, E.; de Rond, L.; Coumans, F. A. W.; Gool, E. L.; Boing, A. N.; Sturk, A.; Nieuwland, R.; van Leeuwen, T. G., Absolute sizing and label-free identification of extracellular vesicles by flow cytometry. Nanomedicine 2018, 14 (3), 801–810.

18. van der Pol, E.; Hoekstra, A. G.; Sturk, A.; Otto, C.; van Leeuwen, T. G.; Nieuwland, R., Optical and non-optical methods for detection and characterization of microparticles and exosomes. J Thromb Haemost 2010, 8 (12), 2596–607.

19. Gasecka, A.; Nieuwland, R.; Budnik, M.; Dignat-George, F.; Eyileten, C.; Harrison, P.; Lacroix, R.; Leroyer, A.; Opolski, G.; Pluta, K.; van der Pol, E.; Postula, M.; Siljander, P.; Siller-Matula, J. M.; Filipiak, K. J., Ticagrelor attenuates the increase of extracellular vesicle concentrations in plasma after acute myocardial infarction compared to clopidogrel. J Thromb Haemost 2020, 18 (3), 609–623.

20. Zhu, S.; Ma, L.; Wang, S.; Chen, C.; Zhang, W.; Yang, L.; Hang, W.; Nolan, J. P.; Wu, L.; Yan, X., Light-scattering detection below the level of single fluorescent molecules for high-resolution characterization of functional nanoparticles. Acs Nano 2014, 8 (10), 10998–1006.

21. Nolan, J. P.; Jones, J. C., Detection of platelet vesicles by flow cytometry. Platelets 2017, 28 (3), 256–262.

22. van der Vlist, E. J.; Nolte-’t Hoen, E. N.; Stoorvogel, W.; Arkesteijn, G. J.; Wauben, M. H., Fluorescent labeling of nano-sized vesicles released by cells and subsequent quantitative and qualitative analysis by high-resolution flow cytometry. Nature protocols 2012, 7 (7), 1311–26.

23. Nolte-’t Hoen, E. N.; van der Vlist, E. J.; Aalberts, M.; Mertens, H. C.; Bosch, B. J.; Bartelink, W.; Mastrobattista, E.; van Gaal, E. V.; Stoorvogel, W.; Arkesteijn, G. J.; Wauben, M. H., Quantitative and qualitative flow cytometric analysis of nanosized cell-derived membrane vesicles. Nanomedicine 2012, 8 (5), 712–20.

24. de Rond, L.; van der Pol, E.; Hau, C. M.; Varga, Z.; Sturk, A.; van Leeuwen, T. G.; Nieuwland, R.; Coumans, F. A. W., Comparison of Generic Fluorescent Markers for Detection of Extracellular Vesicles by Flow Cytometry. Clinical chemistry 2018, 64 (4), 680–689.

25. Welsh, J. A.; Van Der Pol, E.; Arkesteijn, G. J. A.; Bremer, M.; Brisson, A.; Coumans, F.; Dignat-George, F.; Duggan, E.; Ghiran, I.; Giebel, B.; Gorgens, A.; Hendrix, A.; Lacroix, R.; Lannigan, J.; Libregts, S.; Lozano-Andres, E.; Morales-Kastresana, A.; Robert, S.; De Rond, L.; Tertel, T.; Tigges, J.; De Wever, O.; Yan, X.; Nieuwland, R.; Wauben, M. H. M.; Nolan, J. P.; Jones, J. C., MIFlowCyt-EV: a framework for standardized reporting of extracellular vesicle flow cytometry experiments. J Extracell Vesicles 2020, 9 (1), 1713526.

26. Lai, C. P.; Kim, E. Y.; Badr, C. E.; Weissleder, R.; Mempel, T. R.; Tannous, B. A.; Breakefield, X. O., Visualization and tracking of tumour extracellular vesicle delivery and RNA translation using multiplexed reporters. Nat Commun 2015, 6, 7029.

27. Morales-Kastresana, A.; Telford, B.; Musich, T. A.; McKinnon, K.; Clayborne, C.; Braig, Z.; Rosner, A.; Demberg, T.; Watson, D. C.; Karpova, T. S.; Freeman, G. J.; DeKruyff, R. H.; Pavlakis, G. N.; Terabe, M.; Robert-Guroff, M.; Berzofsky, J. A.; Jones, J. C., Labeling Extracellular Vesicles for Nanoscale Flow Cytometry. Sci Rep 2017, 7 (1), 1878.

28. Welsh, J. A.; Horak, P.; Wilkinson, J. S.; Ford, V. J.; Jones, J. C.; Smith, D.; Holloway, J. A.; Englyst, N. A., FCMPASS Software Aids Extracellular Vesicle Light Scatter Standardization. Cytometry A 2020, 97 (6), 569–581.

29. Welsh, J. A.; Jones, J. C.; Tang, V. A., Fluorescence and Light Scatter Calibration Allow Comparisons of Small Particle Data in Standard Units across Different Flow Cytometry Platforms and Detector Settings. Cytometry A 2020, 97 (6), 592–601.

